# Seasonal Variation in the Internet Searches for Cancer Recurrence: An Infodemiological Study

**DOI:** 10.1101/2019.12.12.873984

**Authors:** Xiaoqi Lou, Dingtao Hu, Man Zhang, Qiaomei Xie, Yanfeng Zou, Fang Wang

**Author notes:** Xiaoqi Lou and Dingtao Hu contributed equally to this work and should be considered co-first authors. Corresponding Author Fang Wang, MD, Department of Oncology, The First Affiliated Hospital of Anhui Medical University, 218Jixi Road, Hefei 230022, Anhui, China.

## Abstract

**Background:** While few clinical and epidemiological studies have assessed how seasonality affects cancer recurrence, it has not been studied with the utility of the internet data. In this study, we aim to test whether cancer recurrence presents seasonality on a population level, utilizing internet search query data.

**Methods:** This infodemiological study used Google Trends to find query data for the term “cancer recurrence” from January 01, 2004, to December 31, 2018 in the USA, the UK, Canada, and Australia. Time series seasonal decomposition and the cosinor analysis were used to analyze and describe the seasonal trends for cancer recurrence.

**Results:** A general upward trend in UK and northern hemisphere were observed. Statistically significant seasonal trends on “cancer recurrence” in the USA (*p*=1.33×10^−5^), the UK (*p*=0.012), and northern hemisphere (*p*=5.67×10^−7^) were revealed by cosinor analysis, with a peak in early summer and nadir in early winter. Besides, a seasonal variation was also found in Australia (*p*=2.3×10^−4^), with a peak in late summer and nadir in late winter.

**Conclusions:** The evidence from internet search query data showed a seasonal variation in cancer recurrence, with a peak in early summer(northern hemisphere)/late summer(southern hemisphere). Besides, the relative search volume of “cancer recurrence” appeared a general upward trend in UK and northern hemisphere in recent years.

## Introduction

Cancer, the abnormal proliferation of cells, starts from any organ or tissue and is constituted of tiny cells that unable to form normal functional structures. It is one of the most horrible diseases of the 20th century and spreading further with continuance and rising trend in the 21st century. A quarter of people worldwide is having a risk of cancer during all his or her life (1). Cancer screening and the early detection of cancer offers the opportunity to diagnose cancer early and with an increased opportunity for treatment and curative intent. Nowadays, the improvement of genetic and surveillance technologies, including tumor biomarkers, imaging, and circulating tumor cells and DNA, have contributed to early screening, detection and diagnosis of cancer. These provide more chance for better treatment. Besides, various treatment and management modalities (e.g. surgery, radiation therapy, chemotherapy, immune therapy, targeted therapy, hormone therapy and palliative care) have been applied to oncotherapy. Today, despite advances in early screen, detection, diagnosis, systemic therapies, and patient care, the main reason of death from cancer is due to recurrence that is resistant to therapy. Prolonged monitoring and treatment are warranted for its recurrence. Cancer recurrence could be defined as “after a period of being disease-free, cancer returns” (2,3). It is learnt that people who survive from cancer are at great risk for recurrence of their primary cancer. The distant recurrence risk of the elderly patients who suffered from breast cancer was increasing (4). Recurrent prostate cancer after radical treatment is likely to increase as more people undergo radical treatment (5). Previous studies have also demonstrated that about three quarters of high-risk bladder cancer will experience recurrence, progression, death within 10 years of diagnosis (6). Therefore, it is essential to identify patients at high risk of recurrence or at the state of recurrence to find the characteristics of cancer recurrence, assess curative effect for primary and recurrent cancer, determine the influence of recurrence on death rate, and develop interventions aimed at recurrence.

Recently, web search engines have become pervasive all over the world, so that we can obtain information easily on diverse aspects - from clothing, food, housing, to transportation. Beyond these search interests, however, web-based health condition monitoring and forecasting is getting more and more attention. Google Trends is a web-based tool that displays the input frequency of a specific search word relative to the total search queries. It represents a large, publicly accessible repository of information since the year 2004. Because of its keyword-driven search engines, people all over the world are allowed to obtain online information easily (7,8). After systematically collecting and analyzing, health-related data generated by Google Trends search could be one of the largest databases ever seen in human history, having the possibility to become a significant source of information for healthcare researchers (9). The utility of network data in the field of healthcare research may make up for classical repository of information such as patient investigations and hospital data (9). This method is particularly suited to examine seasonal rhythms of health status and has already been successfully applied to influenza (10), kidney stones (11), and systemic lupus erythematous (12).

To the best of our knowledge, very few clinical and epidemiological studies have assessed how seasonality affects cancer recurrence, and it has not been analysed with the utility of network data. Thus, we performed a study of utilizing network data to validate the hypothesis that cancer recurrence displays seasonal variations.

## Material and methods

### Ethics Statement

This infodemiological study was carried out in conformity to the Declaration of Helsinki, and the Google’s privacy policy (13). It was not necessary for us to require written confirmation of the Medical Ethics Review Committee. The data used in this investigation was publicly available and anonymous data, and involved no personally information such as identity, IP address.

### Google Trends Search and Data Collection

Google Trends (www.google.com/trends) is an online publicly available tracking system, which can automatically record the search volumes of a given term. It is a facility used to assess people’s interest in a user-specified term given a particular location, time period, or category. When people put terms in the Google Trends engine search box, they get data about the frequency of search terms (14). Google Trends allows a random user to compare up to five terms contemporaneously. To make comparisons easier between the terms, it adjusts search data to depict the prevalence of a certain search term within a given time period and geographic region (15). Data points are displayed from 0 to 100 based on a term’s proportion to the total number of searches on all terms (15). Therefore, the data displayed not the actual search times but percentages relative to the total search queries across the given region and time window. The higher the scores, the higher relative search volume (RSV) (14). RSV are easily exported into comma-separated values (CSV) format.

In Jan 18, 2019, Google Trends was mined from inception (01 January 2004) to 31 December 2018 in six different English-speaking countries: the USA, the UK, Canada, Ireland, Australia, and New Zealand. For our study, we searched the query term “cancer recurrence”. Furthermore, to narrow the range of our search, we utilized the “health” category. We were allowed to focus and evaluate interest in the area of health by means of the Health category filter. In this study, the search term ‘cancer recurrence’ was examined utilizing Google Trends. We reasoned that using cancer recurrence as the search term would catch more people who have interested in cancer recurrence. This supposition was validated by the fact that ‘cancer recurrence’ on Google Trends revealed a higher RSV than all other combinations of these terms (e.g. cancer, tumor, carcinoma, neoplasm, recurrence, relapse, recur, recrudescence, recrudesce, reappearance, reappear). The USA, the UK, Canada and Ireland were chosen to represent the native English-speaking countries in the Northern hemisphere, and Ireland and Australia were representatives of Southern hemisphere accordingly. However, two countries (Ireland, New Zealand) were ruled out in the analysis. The reasons excluding Ireland were limited Google Trends data and near-zero search volume more than 12 months in a row, while the search volume in New Zealand was not available because of the finite population. Hence, the representative countries of the Northern hemisphere were the USA, the UK and Canada, while Australia represents the Southern hemisphere. We added up the monthly search volume of the USA, the UK and Canada to form the monthly search volume of northern hemisphere. This strategy allowed us to evaluate the seasonality of search queries that should be out of phase by approximately 6 months between these hemispheres, as presented in earlier studies (16,17,18).

The monthly data used in this study were gathered by manually organizing every data point on a specific Google Trends graph. For the reproducibility and completeness of our search, the CSV data of each data point were recorded at the time of collection.

### Statistical Analysis

Time series seasonal decomposition analysis was applied to represent the trend and seasonality in a time series. All time series data were decomposed into a trend component, a seasonal component and a remainder component. This analysis was designed to identify seasonal variations in cancer recurrence in the study period (from 2004 to 2018).

Similar to existing studies analyzing the seasonal trends of health status utilizing network data (19,20,21), the cosinor analysis was applied to validate the hypothesis that the RSV of cancer recurrence displays significant seasonality during the period of study. The cosinor analysis and the software used for execution are given a detailed description elsewhere (22). In brief, cosinor analysis depends on a sinusoid

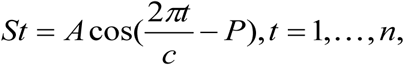

in which *A* is the amplitude of the sinusoid, *c* is the length of the seasonal cycle (set at 12 for monthly data), *t* is the time of each data point, *P* is the phase, and *n* is the total number of data points (23). The amplitude states the size of seasonality, and the phase states the time of seasonal spike. As mentioned before, *n*=180. Given that the cosinor model is part of a generalized linear model, we could calculate the statistical significance of any seasonal trends. Since the seasonal component of the sinusoid is consist of sine and cosine, all represented *p* values are the half of the original *p* value for the purpose of correcting for multiple comparisons (19,20). Namely, the adjusted statistical significance p value is *p* < 0.025. There are two p values in the cosinor model, one for the sine and the other for the cosine. And then both of the *p* values are checked to identify the statistical significance based upon the adjusted statistical significance *p* value. In the end, either sine or cosine *p* value was showed (22). Additionally, a time series plot was executed to highlight the consistent seasonality in cancer recurrence.

Seasonal and trend decomposition plots, cosinor analyses and time series plots were executed with the use of the ‘season’ package (22) in R version 3.5.1.

## Results

### Analyses of Systematic Seasonal Variations and Trends for the Relative Search Volume

Analyses of seasonal variations and trends for cancer recurrence in the USA, UK, Canada, Australia, and northern hemisphere were presented in Fig. 1. It was obvious to see that the variation of the RSV of cancer recurrence in the four countries and northern hemisphere mentioned above showed the cyclicity with 12 months being a circle, and showed a clear seasonal patterns from 2004 to 2018. Meanwhile, we detected a general upward trend in UK and northern hemisphere, and the general upward trend of the northern hemisphere mainly resulted from the increasing RSV of UK.

**Fig 1.**
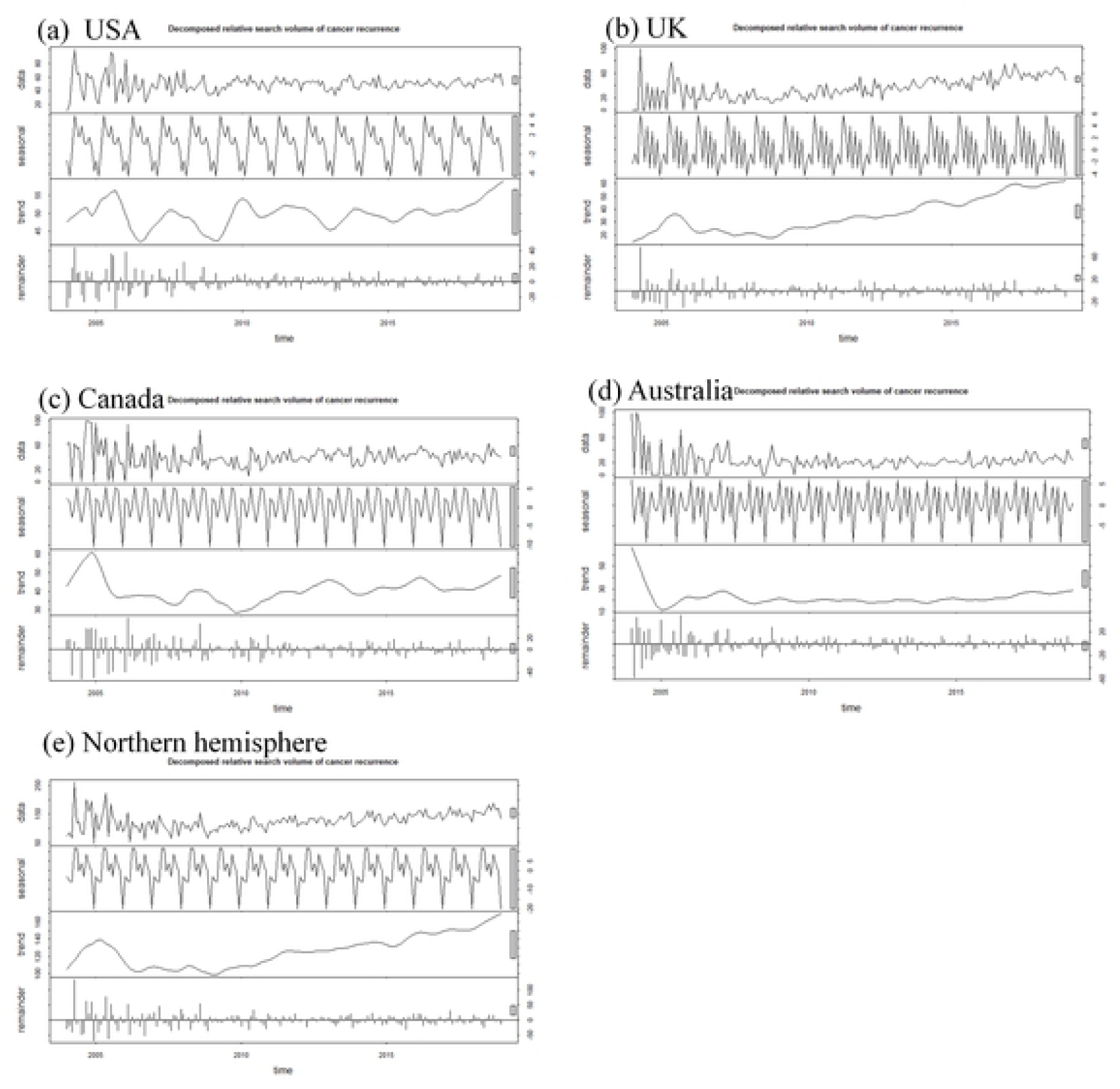
Seasonal and trend decomposition for the relative search volume of “cancer recurrence” from January 01, 2004, to December 31, 2018

### Cosinor Analysis for the Relative Search Volume

The results of the cosinor analysis were shown in Table 1, and the plots of cosinor models were displayed in Fig. 2. The search query data for the United States, the United Kingdom, Canada, northern hemisphere, and Australia showed clear peaks and nadirs, which could be assessed by visual inspection. Cosinor analysis largely validated this with statistically significant seasonal trends on ‘cancer recurrence’ in the USA (amplitude [*A*] = 3.54, phase month [*P*] = 6.4, low point month [*L*] = 12.4, p=1.33×10^−5^), the UK (*A*=1.73, *P*=6.3, *L*=12.3, *p*=0.012), northern hemisphere (*A* = 6.44, *P* = 6.4, *L* = 12.4, *p*=5.67×10^−7^), and Australia (*A* = 2.22, *P* = 2.0, *L* = 8.0, *p*=2.3×10^−4^). Obviously, the peak for both countries was in the summer (June or early summer for the northern hemisphere countries; February or late summer for the southern hemisphere country) and nadir in the winter (December or early winter for the northern hemisphere countries; August or late winter for the southern hemisphere country). The peaks (early summer/late summer) and nadirs (early winter/late winter) were out of phase by approximately 6 months in the northern hemisphere countries compared with the southern hemisphere country. Meanwhile, as shown in Table 1, in USA, UK, and Canada, the top ranking search cancer was breast cancer consistently. Therefore, the seasonal patterns of cancer recurrence may be embodied in breast cancer.

**Table 1.** The seasonal variation in the relative search volume of “cancer recurrence”.

| Country | Amplitude | Phase month | Low point month | *P*-value | Top ranking search cancer |
| --- | --- | --- | --- | --- | --- |
| USA | 3.54 | 6.4 | 12.4 | $1.33\times10^{-5}$ | Breast cancer |
| UK | 1.73 | 6.3 | 12.3 | 0.012 | Breast cancer |
| Canada | 1.19 | 6.4 | 12.4 | 0.100 | Breast cancer |
| Northern hemisphere | 6.44 | 6.4 | 12.4 | $5.67\times10^{-7}$ | — |
| Australia | 2.22 | 2.0 | 8.0 | $2.3\times10^{-4}$ | No data |

**Fig 2.**
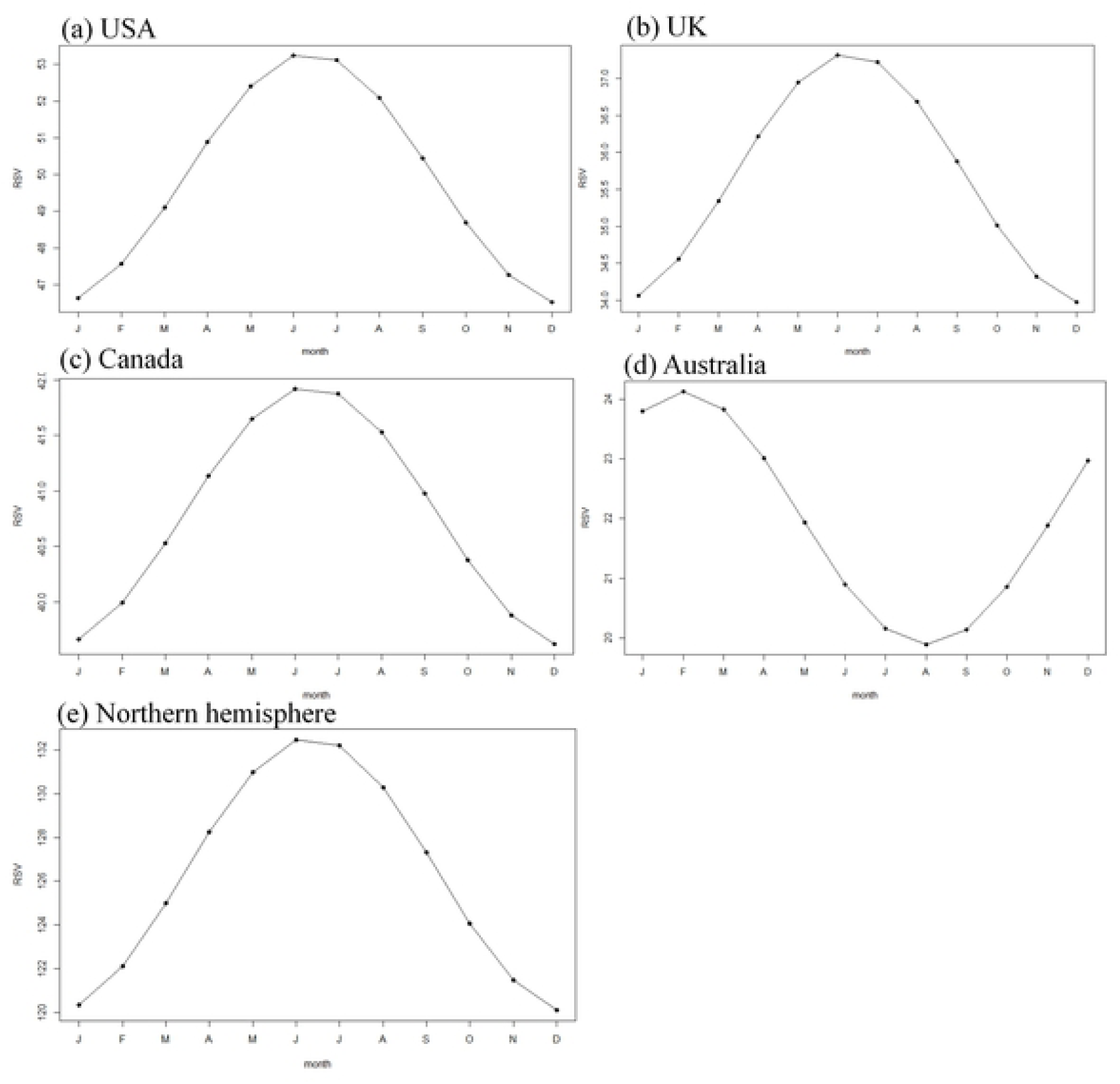
The plots of cosinor models for the seasonal variation in the relative search volume of “cancer recurrence”

According to time series plots shown in Fig. 3, Google searches for the term ‘cancer recurrence’ showed a general upward trend throughout the study period (from 2004 to 2018) in UK, Australia and northern hemisphere, and others countries (the USA, Canada) in northern hemisphere didn’t show such trend.

**Fig 3.**
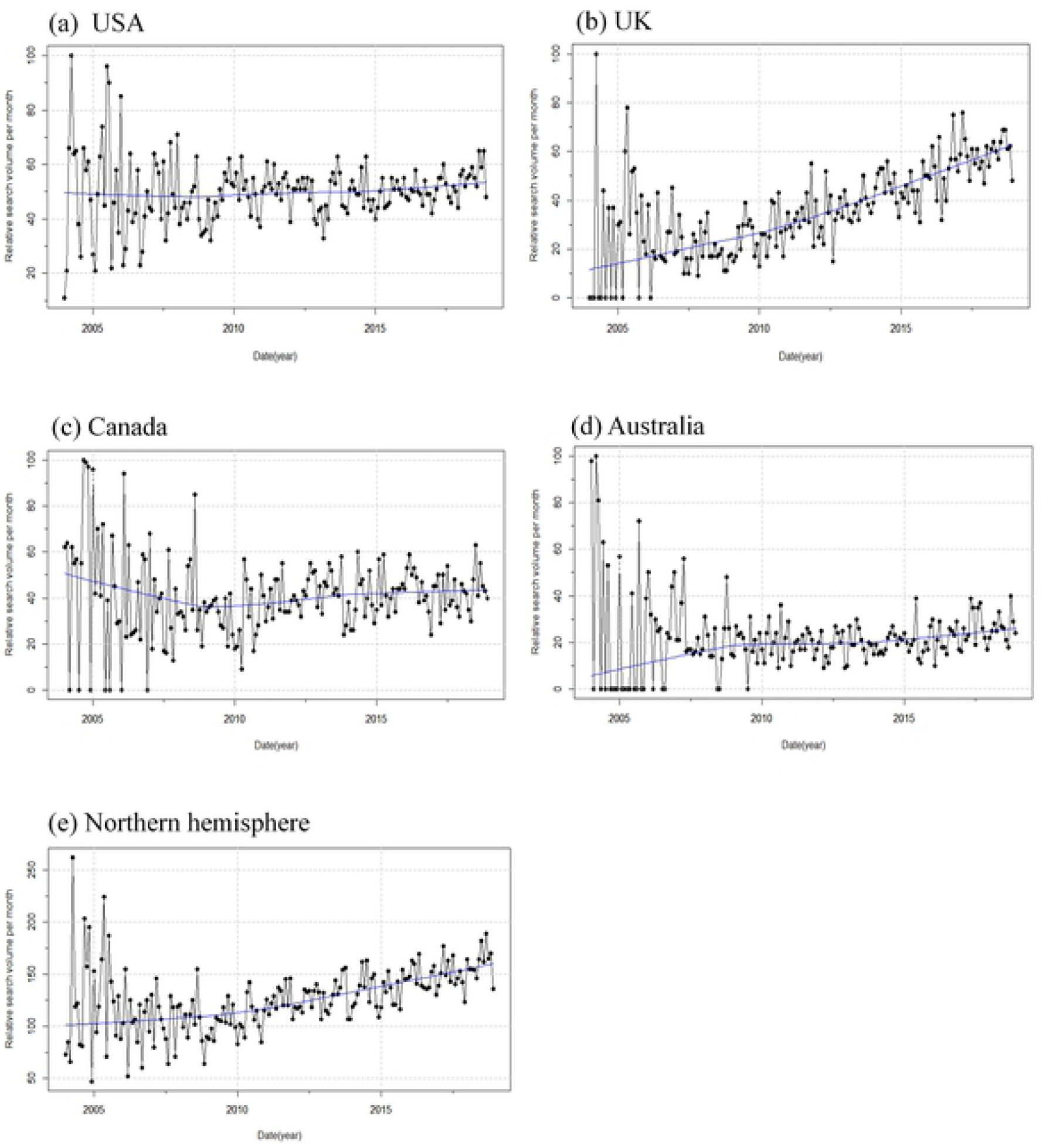
Time series plots for the relative search volume of “cancer recurrence” from January 01, 2004, to December 31, 2018

Given the above, there were mutual authentication relationship between analyses of systematic seasonal variations and trends and cosinor analysis, both of them showing a general upward trend over the study period (from 2004 to 2018) in UK and northern hemisphere. Whereas, compared with time series seasonal decomposition analysis, Cosinor analysis provided more detailed information to us, and it elaborated the amplitude, peak month, low point month and *p* value.

## Discussion

This study analyzed the seasonality of “cancer recurrence” across several countries, utilizing internet search query data. Results showed a significant seasonal trend for ‘cancer recurrence’ in the USA, UK, Australia, and northern hemisphere, with a peak in early summer/late summer and nadir in early winter/late winter. The peaks (early summer/late summer) and nadirs (early winter/late winter) were out of phase by approximately 6 months in the northern hemisphere countries compared with the southern hemisphere country. Besides, we also detected a general upward trend in UK and northern hemisphere, and the general upward trend of northern hemisphere mainly resulted from the increasing relative search volume of UK. As our results suggested, in USA, UK, and Canada, the top ranking search cancer is breast cancer, so the seasonal variation that we found in cancer recurrence may be primarily due to breast cancer.

So far, this is the first study exploring the seasonality of cancer recurrence. Though, the mechanisms underlying the seasonal trends of cancer recurrence detected in our study cannot be assessed, there are several possible factors (e.g., diet, physical exercise, smoke, alcohol, air pollution, infections, immunity and endocrine dyscrasia) that may associated with the seasonality of cancer recurrence. Firstly, the immune functional activity of neutrophils was increased in summer, which was associated with poor recurrence-free survival (24,25). Secondly, as our results suggested, in USA, UK, and Canada, the top ranking search cancer is breast cancer, so the seasonal variation that we found in cancer recurrence may be primarily due to breast cancer. Previous study has indicated that melatonin, a kind of indole hormone produced mainly by the pineal body, could result in growth inhibition of all three estrogen-responsive human breast tumor cell lines (MCF-7, T47D, ZR-75-1) (26). Therefore, an increase in melatonin levels is not conducive to recurrence of breast cancer. However, melatonin levels are lower in summer than in winter (27,28). This provides another piece of proof for the unanimous seasonal trends in cancer recurrence. Furthermore, alcohol use in spring and summer is higher than in fall and winter (29,30), which may lead to the increasing risk of breast cancer recurrence to some extent (31). Thirdly, alcoholism, which prevail in spring and summer (29,30), promotes the risk of HCC recurrence amongst non-B or non-C (NBNC) hepato-cellular carcinoma (HCC) patients (32,33). Fourthly, the peak of urinary tract infection (UTI) is in summer (34), and Kim et al. have demonstrated that pyuria is a risk factor for bladder cancer recurrence during short-term follow-up after urethral and bladder tumor resection (TURBT) (35). Lastly, compared with winter, the increase of PSA in the body in summer is more common, and the increase of PSA indicates the recurrence of prostate cancer after the initial treatment, leading to a higher recurrence rate of prostate cancer in summer (36,37,38).

According to time series seasonal decomposition and the cosinor analysis, the relative search volume of cancer recurrence revealed a general upward trend in UK and northern hemisphere, and the general upward trend of the northern hemisphere mainly resulted from the increasing relative search volume of UK. Over the past decade, the morbidity rates of all cancers combined have increased by 7% in the UK (39), may resulting in high relative search volume of cancer recurrence, and then causing the upward trend. Consistently demonstrated by Eurocare and the International Cancer Benchmarking Partnership (ICBP), the survival rates of cancer patients in the UK were greatly lower than in other developed countries, which may be explained by the increasing cancer recurrence rates.

There are several advantages in our study including the substantial and detailed mass of data, the long-term of observation, the wide areas of coverage (including countries on both sides of the equator), and the lack of reporting and observer bias. Despite the strength mentioned above, there are inherent disadvantages in our study remain to be discussed. Firstly, the data do not supply the demographics of people who participate in the web search query, and thus significant inter-individual differences (e.g., age, gender, etc.) that may be associated with the likelihood of utilizing net-based health information cannot be evaluated. Therefore, this study only in favor of population-level seasonal variations of cancer recurrence. Secondly, inherent to the use of web-based query data in the study of disease is the assumption that such queries represent the prevalence of disease and/or the severeness of disease, the veracity of which cannot be evaluated. Thirdly, the reason why selection bias exist in the process of statistical analysis is that we used data from a single search engine, Google. Nevertheless, this risk is alleviated by the truth that Google makes up more than 65% of all web searches worldwide (40).

In conclusion, Internet queries for cancer recurrence display significant seasonality, with a peak in the summer, and nadir in the winter. Further studies are needed to elucidate the mechanisms of the seasonal variations in cancer recurrence in the population. These seasonal effects may exert influence on clinical practice. We can take many measures during the summer to decrease the recurrence rates, such as increase the frequency of reexamination, limit alcohol intake, and reinforce prevention measures for UTI.

## Acknoledgements

The authors acknowledge the National Natural Science Foundation of China (81602115,81872683,81673254), the Foundation of Supporting Program for the Excellent Young Faculties in University of Anhui Province in China (gxyq2019012), Grants for Scientific Research of BSKY from the First Affiliated Hospital of Anhui Medical University and Grants for Outstanding Youth from the First Affiliated Hospital of Anhui Medical University.

## Author contributions

**Conceptualization:** Fang Wang, Yanfeng Zou

**Formal analysis:** Fang Wang, Yanfeng Zou, Xiaoqi Lou, Dingtao H

**Funding acquisition:** Fang Wang

**Investigation:** Fang Wang, Xiaoqi Lou, Dingtao H

**Methodology:** Fang Wang, Xiaoqi Lou, Dingtao H

**Writing-original draft:** Xiaoqi Lou, Dingtao Hu

**Writing-review & editing:** Xiaoqi Lou, Dingtao Hu, Man Zhang, Qiaomei Xie

